# Explainable machine learning relates histological to genomic pathology

**DOI:** 10.64898/2026.08.03.742582

**Authors:** John Connelly, Barbara Hernando, Juliet Luft, Craig J. Anderson, Peter Bankhead, Frances Connor, Stuart Aitken, Liver Cancer Evolution Consortium, Colin A. Semple, Paul Flicek, Duncan T. Odom, Martin S. Taylor, Sarah J. Aitken

**Author notes:** Co-first author (John Connelly, Barbara Hernando). **Correspondence** Correspondence and requests should be addressed to Martin S. Taylor and Sarah J. Aitken. A list of authors and their affiliations appears at the end of the paper.

## Abstract

**Background & Aims:** Haematoxylin and eosin (H&E) staining remains the diagnostic gold standard for solid cancers, including hepatocellular carcinoma, and is increasingly complemented by genomic profiling for precision medicine. Inferring genomic alterations directly from H&E images could streamline testing, but heterogeneity and biases in human training data limit interpretation of genotype-phenotype associations. Here, we aimed to relate histologic to genomic pathology to provide biological explainability for mutation prediction models and assess the impact of germline variation on model performance.

**Methods:** We analysed 597 murine liver tumours with matched whole-genome sequencing and histopathology (163,835 image tiles; 22.9 million nuclei). Our controlled *in vivo* design accounted for germline variation, biological sex, and causal mutagen (N-diethylnitrosamine), removing confounding factors present in human cohorts. We trained and evaluated deep learning and supervised machine learning models to predict germline variation and cancer driver alterations from H&E.

**Results:** Modelling accurately predicted germline and somatic alterations from histology, at both locus-specific and genome-wide scales. Quantitative image analysis revealed an unexpected association between *Egfr* driver mutations and hepatic steatosis, linking genotype to an interpretable morphological phenotype. While model performance declined when applied to tumours from unrepresented genetic backgrounds, this limitation was biologically informative, revealing strain-dependent differences in tumour evolution, notably the prevalence of whole-genome duplication.

**Conclusions:** Machine learning integration of histological and genomic pathology enables accurate, interpretable inference of genetic alterations from H&E, potentially reducing reliance on costly ancillary molecular assays. Our predictions are supported by human-interpretable biological features, addressing concerns around “black-box” technologies. However, caution is required when applying such methods to samples with a genetic background that, even if closely related, is beyond the genetic horizon of training data.

## Introduction

Large-scale cancer sequencing studies have accelerated our understanding of tumour biology by identifying clinically important molecular biomarkers^1–4^. Cancer genomes, like the tissue morphology of a biopsy or resection, give a static snapshot of a dynamic evolutionary disease process. Tissue morphology therefore provides a phenotypic read-out of the underlying tumour genome, reflecting the cumulative effects of mutational, repair, and selective processes that shape tumour evolution.

Histopathology is the gold-standard for the diagnosis of solid organ cancers, including hepatocellular carcinoma (HCC). By interpreting the cellular morphology of haematoxylin and eosin (H&E) stained tissue sections, pathologists determine the diagnosis, grade, and stage of disease^5,6^, which also provides prognostic information to guide treatment decisions. The addition of molecular pathology (e.g. immunohistochemistry (IHC) and fluorescent *in situ* hybridisation (FISH)) and genomic pathology assays (e.g. targeted gene panels and exome, genome, or transcriptome sequencing) can assist differential diagnosis, tumour subtyping, or provide predictive information about actionable molecular pathways, driver mutations or signatures^7–11^. For example, the differential diagnosis of HCC is often assisted by IHC for heat shock protein-70 (HSP-70), glypican-3 (GPC-3), and glutamine synthetase (GS)^12,13^, and expression of cytokeratin 19 (CK19) or Ki67 are associated with poor prognosis^14,15^. However, despite the promise of ancillary molecular biomarkers, their clinical implementation is often limited by practical constraints. These include physical factors (e.g. differences in tissue processing^16^, paucity of availability tissue, or unpredictable cold chains), temporal delays (e.g. real-world implementation of even rapid reflex testing can delay diagnosis, in turn impacting tumour board decisions and patient treatment), and financial barriers (e.g. limited resources or lack of reimbursement). Consequently, computational pathology approaches that extract actionable molecular information directly from morphology (i.e. “image biomarkers” derived from digitised H&E slides) would offer an alternative with substantial clinical benefit.

Deep learning algorithms have already demonstrated varying success at: (i) replicating some diagnostic capabilities of the pathologist in classifying neoplasia^17–22^, (ii) inferring ‘invisible’ information that is not discernible to the human expert, but that is associated with patient survival^23–26^, and (iii) predicting point mutations, copy number variants, and gene expression signatures from minimally-processed human pan-cancer datasets^27,28^. These studies provide evidence that deep learning may be able to triage, or even eventually replace, genetic testing for precision oncology^29,30^. However, despite high predictive performance, deep learning is frequently criticised as lacking explainability, interpretability, and human-intuition^31^; this is an important barrier to widespread adoption by practising pathologists and healthcare regulators^21^ and poses a major challenge to moving morpho-molecular inference models into the clinical domain^32^.

Human cancer cohorts exhibit extensive genomic complexity and heterogeneity, reflecting diversity of ancestry backgrounds, disease aetiology, and environmental and/or therapeutic exposures. Heterogeneity in tumour pathology poses a challenge for the training, evaluation, and implementation of AI models which is exacerbated by sparse metadata, and further biased when cohorts are not fully representative of patient groups or the wider population. For example, there are known differences in cancer incidence^33^, histological subtypes^34–36^, and molecular landscape^37–43^ between ancestry groups, but it remains unknown whether this is also reflected in cellular morphology.

To address this limitation, we have applied deep learning approaches to an experimental cohort of murine liver tumours across divergent genetic backgrounds. Briefly, we re-ran tumour evolution hundreds of times to generate 597 chemically-induced liver tumours from four inbred strains of mice and analysed their whole genome sequencing^44,45^ and matched digitised histopathology whole slide images (WSIs) (**Methods; Fig. 1A**). Our prior genome-wide analyses revealed the majority of these histologically near-identical tumours displayed genome-wide strand asymmetry of mutations^44,45^. This unique, paired histological and genomic pathology dataset provides the opportunity to study phenotype-genotype associations using computational pathology and machine learning, while controlling major covariates including germline genetic variation, environmental exposure, and tumour cell-of-origin. A similarly controlled experimental design is not possible in human cohorts. We sought to address both methodological and biological questions. Specifically, are deep transfer learning methods appropriate to predict specific molecular/genetic changes from H&E stained tissue sections - and is model prediction impacted by genetic background? Additionally, can supervised machine learning be used to generate biological hypotheses about the underlying molecular mechanisms that drive these predictions and identify human-intuitive features to explain phenotype-genotype associations? By combining these orthogonal approaches, we aimed to identify quantitative, interpretable (but unbiased, machine-detected) information that provides biological explainability and overcomes the limitations of “black-box” AI models.

**Fig.1.**
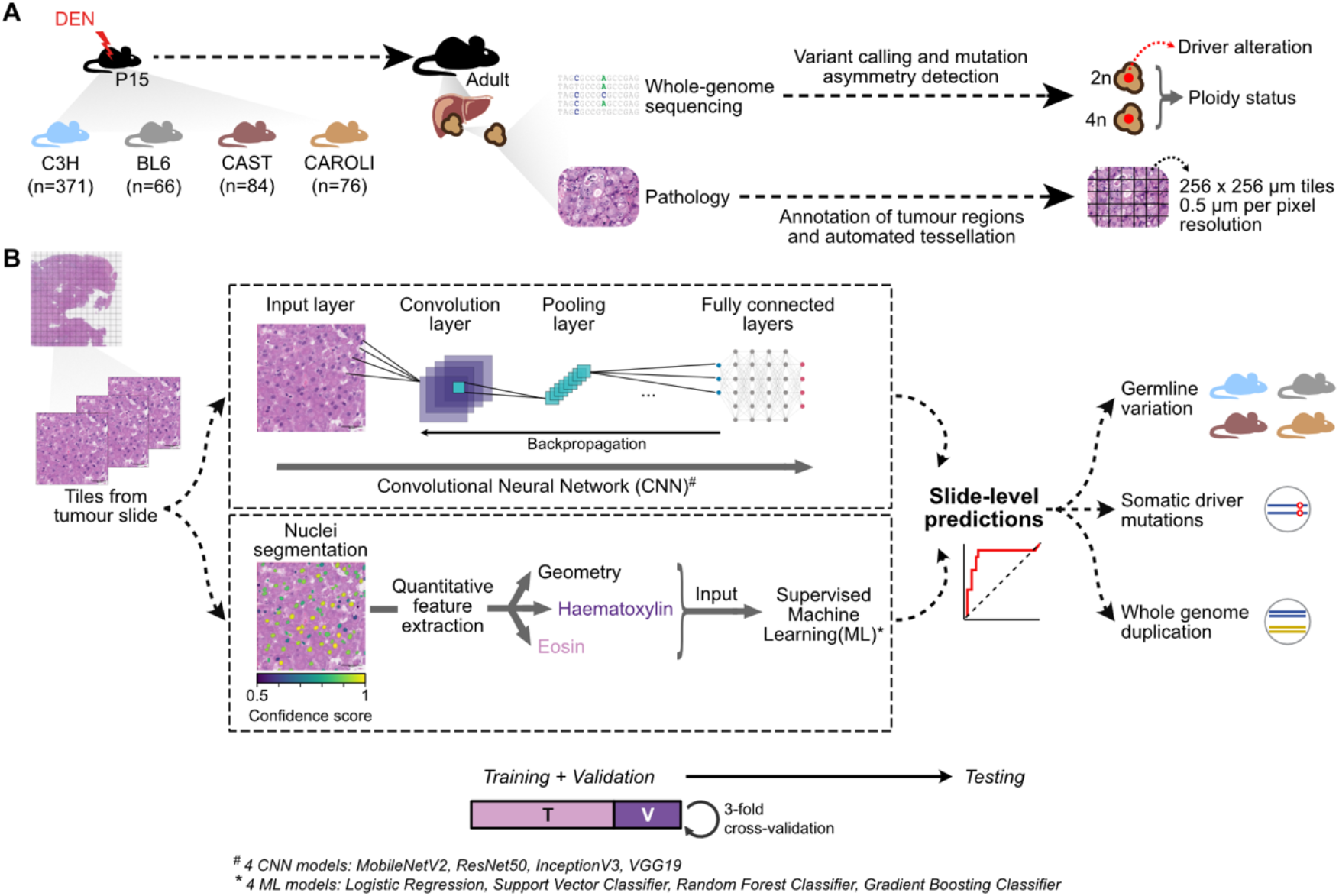
Overview of experimental and computational approaches. **(A)** Liver tumours (n=597) were chemically-induced using diethylnitrosamine (DEN) in four evolutionarily divergent strains of mice (C3H, BL6, CAST, and CAROLI). Tumours were bisected and analysed using whole genome sequencing (WGS) and computational pathology of digitised H&E slides. Driver alterations, ploidy status, and tessellated tumour tiles were used as input for phenotype-genotype modelling. **(B)** Weakly supervised deep transfer learning pipelines based on Convolutional Neural Networks (MobileNetV2, ResNet50, InceptionV3, VGG19) were applied to H&E tessellated tiles (n=163,835). Supervised machine learning approaches (Logistic Regression, Support Vector Classifier, Random Forest Classifier, Gradient Boosting Classifier) were applied to quantified nuclear features (n=22,879,653; geometry and colourimetry). These models were then used to make slide-level predictions (AUC-ROC) of genetic alterations: germline variation, somatic driver mutations, and whole genome duplication.

## Materials and Methods

The data analysed in the manuscript were generated for two parallel studies^44,45^ and are publicly available. Whole slide images (WSIs) are available from BioStudies: S-BSST383 ^44^ and S- BSST384 ^45^. Whole genome sequencing FASTQ files are available in the European Nucleotide Archive: PRJEB37808 ^44^ and PRJEB15138 ^45^.

Full details of the experimental methodologies are provided in those manuscripts, and briefly summarised below to provide pertinent details relevant to the histopathology image analysis and genomic analyses.

### Mouse colony management

Animal experimentation was carried out in accordance with the Animals (Scientific Procedures) Act 1986 (United Kingdom) and with the approval of the Cancer Research UK Cambridge Institute Animal Welfare and Ethical Review Body (AWERB).

The following mouse strains and species were used: *Mus musculus domesticus* C3H/HeOuJ (hereafter referred to as C3H mice) and C57BL/6J (BL6), *Mus musculus castaneus* CAST/EiJ (CAST), and *Mus caroli* CAROLI/EiJ (CAROLI). For simplicity, these strains, subspecies and species are herein referred to as ‘strains’.

Half of the BL6 liver tumours were induced in mice with hepatocyte-specific *Ctcf* hemizygosity (*Ctcf*^+/-^); the remaining BL6 tumours were induced in their wild-type (*Ctcf*^+/+^) littermates. Male mice with whole-body *Ctcf* hemizygosity (the *Ctcf* allele has loxP sites inserted at exon 3 and exon 12; full details in^46^) were crossed with female mice expressing Cre recombinase under the control of the albumin promoter. Since albumin is highly and uniquely expressed by hepatocytes, this cross results in liver-specific knock-out of one *Ctcf* allele using the Cre/loxP system; all non-hepatocytes in the mouse retain two copies of *Ctcf*. Littermates had the expected 1:1 Mendelian ratio of genotypes^46^.

### Chemical model of hepatocarcinogenesis

15-day-old (P15) male mice of all strains were treated with a single intraperitoneal (IP) injection of N-Nitrosodiethylamine (DEN; Sigma-Aldrich N0258; 20 mg/kg body weight) diluted in 0.85% saline. Liver tumour samples were collected from DEN-treated mice 25 weeks (C3H), 36 weeks (BL6), 38 weeks (CAST), or 78 weeks (CAROLI) after treatment; pilot data indicated that 100% of surviving DEN-treated mice would develop tumours by these timepoints. Liver tumours of sufficient size (≥2 mm diameter) were bisected; one half was flash frozen in liquid nitrogen and stored at -80°C for DNA and RNA isolation, and the other half was processed for histology.

All mice included in the original study were male which controls for metabolic and cancer susceptibility differences between biological sexes; an equivalent retrospective cohort of female mice was not available to analyse in parallel.

### Histopathology and whole slide image acquisition

Tissue samples for histology were fixed in 10% neutral buffered formalin for 24 h, transferred to 70% ethanol, machine processed (Leica ASP300 Tissue Processor; Leica, Wetzlar, Germany), and paraffin embedded. All formalin-fixed paraffin-embedded (FFPE) sections were 3 μm in thickness. Haematoxylin and eosin (H&E) staining was performed using the automated Leica ST5020; mounting was performed on the Leica CV5030. Tissue sections were digitised using the Aperio XT system (Leica Biosystems) at 20x resolution.

Steatosis was quantified in C3H liver tumours by analysing H&E stained WSIs (Aperio SVS format) using Visiopharm (version 2020.08; application 10119). Tissue detection was based on H&E, minimum droplet size for inclusion was set to 5μm^2^, and the small/large droplet threshold was set to 20μm^2^.

### Tumour diagnosis

H&E sections of liver tumours were blinded and assessed twice by a board certified histopathologist (S.J.A) according to the International Harmonization of Nomenclature and Diagnostic Criteria for Lesions in Rats and Mice (INHAND) guidelines^47^.

### Tumour inclusion criteria

Tumours which met the following histological criteria were selected for whole genome sequencing: (i) diagnosis of dysplastic nodule (DN), (ii) homogenous tumour morphology, (iii) tumour cell percentage >70%, and (iv) adequate tissue for DNA extraction. Those with extensive necrosis, mixed tumour types, a nodule-in-nodule appearance (indicative of a hepatocellular carcinoma (HCC) arising within a DN), or contamination by normal liver tissue were excluded. For tumours with multiple transections, only a single slice was included.

### Whole genome sequencing, alignment, and variant calling

Whole genome sequencing (WGS) libraries were generated using TruSeq PCR-free Library Prep Kit (Illumina 20015963) and sequenced on a HiSeq X Ten (Illumina) to produce paired-end 150 bp reads (minimum of 40x coverage). Sequencing reads were aligned to their respective genome assemblies from Ensembl (v.91)^48^ (BL6, GRCm38, GenBank: GCF_000001635.26; C3H, C3H_HeJ_v1, GenBank: GCA_001632575.1; CAST, CAST_EiJ_v1, GenBank: GCA_001624445.1; CAROLI, CAROLI_EiJ_v1.1, GenBank: GCF_900094665.2) using bwa-mem (v.0.7.12)^49^. Variant calling and filtering were performed as previously reported^44,45^. Three driver discovery approaches were implemented: OncodriveFML (v.2.2.0)^50^, OncodriveCLUSTL (v.1.1.3)^51^, and dNdScv (v0.1.0)^52^. Tumours with either multiple MAPK driver mutations or no identified driver mutation were excluded from all downstream analyses.

### Classification of mutation strand asymmetry

Genomic segmentation of mutational asymmetry was performed as previously reported44,45. Mutational strand asymmetry was scored for each genomic segment using the relative difference metric S=(W-C)/(W+C) where W is the rate of mutations from T on the forward (plus) strand of the reference genome and C the rate of mutations from A on the plus strand. Genomic segments were classified as symmetric (abs(S) < 0.2, where ‘abs(S)’ is the absolute value of the ‘S’ parameter), asymmetric (abs(S) > 0.8), or intermediate (abs(S) ≥ 0.2 and ≤ 0.8). For each tumour, the proportion of the genome belonging to each class was calculated. For assigning dichotomous genome symmetry status, samples with 0% asymmetric autosomes and <25% intermediate autosomes were defined as non-asymmetric (i.e. symmetric)-genome, otherwise samples were defined as asymmetric-genome.

### Image preprocessing

597 whole slide images of liver tumours were manually annotated by two pathologists (J.C. and S.J.A) using QuPath (v.0.2.2)^53^ to exclude adjacent normal parenchyma, cystic spaces, processing artefacts, and white space. Annotated tumour regions were tessellated into fixed-size, non-overlapping square 256 µm x 256 µm tiles (0.5 µm per pixel resolution) using Groovy in QuPath, and written as TIFFs for nuclear segmentation and morphometry quantification, and as PNGs for deep transfer learning. As expected, the number of tiles is strongly correlated with the size of the annotated tumour (**Supplementary Fig. 1A**). To account for staining differences and batch effects, PNGs images were normalised using the Macenko method^54^ as implemented in torchstain (v.1.2.0)^55,56^.

### Nuclear segmentation and feature quantification

Epithelioid nuclei were segmented using a pre-trained StarDist^89^ model (he_heavy_augment.zip) downloaded from https://github.com/stardist/stardist-imagej/tree/master/src/main/resources/models/2D and an inference instance was deployed using Groovy across the tiles in QuPath, built from source with TensorFlow^57^, with a minimum detection threshold of 0.5. Following stain deconvolution^58^ using default vectors, fifteen quantitative nuclear features were measured per nuclear object, reflecting geometry (size and shape) and colourimetry (histochemical staining intensity of haematoxylin and eosin). Optical density is a measure of the absorbance of light, proportional to stain intensity; the greater the amount of stain present, the greater the optical density. The object-level measurements were subsequently summarised by computing the median, median, standard deviation, interquartile range, and kurtosis for each tile and WSI. Nuclear volume was calculated using the median nuclear diameter per slide.

Filtering was applied to exclude: tiles with a nucleus count ≤1st percentile; nuclei with an area ≥99.9th percentile, or circularity ≤0.01st percentile or non-calcuable.

One tile per WSI was selected randomly for manual validation of type I and type II error in the identification of all epithelioid nuclei. This demonstrated that average precision and detection probability were not systematically biased by strain, driver status, or genome symmetry (**Supplementary Fig. 1B-D**).

### Deep transfer learning from image tiles

Tumours were partitioned into training (70%) and test (30%) cohorts using SciKit-Learn (v.1.0.2)^59^, partitioning was done at slide or batch level which ensures that all tiles from the same tumour are within the same training (or test) set, in turn avoiding data leakage. Within the training set, tiles in the majority class were randomly downsampled to ensure parity with the minority class for all experiments (with the exception of Ctcf).

Deep learning was performed using TensorFlow (v.2.9.1)^57^ as described in the library documentation^60^. Four Convolutional Neural Networks (CNNs) models (MobileNetV2 ^61^, ResNet50 ^62^, InceptionV3 ^63^, and VGG19 ^64^) were downloaded from the TensorFlow API. These models were initially re-trained with a convolutional base frozen to ImageNet weights, then fine-tuned with the deepest one-third of layers unfrozen. PNG format images were used for deep transfer learning. Images were parsed through training in batches of 20 using the ‘image_dataset_from_directory’ function within TensorFlow with both prefetching and autotuning. Images were sized according to the bespoke requirements of each CNN and the requisite preprocessing function provided within TensorFlow was applied for each CNN.

Training was performed with Graphics Processing Unit (GPU) acceleration on an NVIDIA RTX A5000 (driver: 470.141.03) with CUDA Toolkit (v.11.3.1) and cuDNN (v.11.3.1)^65^. An Adam optimiser, with a learning rate of 0.001 up to 10 epochs was used for initial frozen-base training, then an RMSprop optimiser with a learning rate of 0.0001 up to a further 10 epochs was used for fine-tuning. Cross-entropy (binary or categorical as appropriate) was used as the loss function and accuracy (binary or categorical) as the evaluation metric for both frozen base and fine tuning. The models were regularised with 20% dropout and early stopping checkpoints. Formal hyperparameter search was not performed for these models. Fitted models were persisted in Hierarchical Data Format (version 5, HDF5) format.

For task inference, tile-level prediction probabilities were initially generated. The tile-level values of an individual WSI (tumour) were then used to generate a ROC curve and derive performance metrics per tumour, including AUC, precision, recall, balanced accuracy and F1 scores. Confidence intervals were estimated from 1,000 bootstrap samples of tiles per WSI.

### Supervised modelling of quantitative nuclear features

Tumours were partitioned for training and testing as above (70/30), either at batch level or at WSI level. Tiles were treated as the instance unit for supervised model training. The training set was optimised using the SMOTEEN (minority over-sampling and nearest neighbours cleaning) method from imbalanced-learn (v.0.7.0)^66^.

Four learning algorithms (logistic regression, support vector classifier, random forest classifier, and gradient boosting classifier) were trained using SciKit-Learn (v.1.0.2). The logistic regression models were fitted with elastic net regularisation. The support vector classifier was fitted with an RBF kernel. Model training was implemented by combining 3-fold cross validation with successive halving grid search. Model objects were fitted using SciKit-Learn’s pipeline class and included feature normalisation by Z-scoring using the StandardScaler function. For the logistic regression and support vector classifier, dimensionality reduction was performed by principal component analysis, with the number of principal components selected by the hyperparameter search. A hyperparameter search dictionary was defined for each model pipeline. Training was accelerated by multi-core processing using joblib^67^ parallelisation.

Fitted models were persisted joblib objects. For task inference, tile-level prediction probabilities were initially generated. The tile-level values of an individual WSI (tumour) were then used to generate a ROC curve and derive performance metrics per tumour, including AUC, precision, recall, balanced accuracy and F1 scores. Confidence intervals were estimated from 1,000 bootstrap samples of tiles per WSI.

### Global explanations of supervised models

SHAP (v.0.40.0)^68^ was used to compute Shapley values, with respect to the positive class, for each of the random forest classifiers fitted to the full feature space without dimensionality reduction. Global model explainability was visualised using the ‘summary_plot’ function of the SHAP package, with the number of features displayed limited to 10.

### Computational analysis environment

The majority of the codebase for image analysis and machine learning was implemented in Python 3 and Jupyter-Lab (v.3.2.1). General data manipulation and numerical tasks were performed using Pandas (v.1.3.4)^69^ and Numpy (v.1.20.3)^70^. Statistical testing was performed using statannot (v.0.2.3)^71^ and scipy (v.1.7.3)^72^. Principal component analysis was performed using Sci-Kit Learn (v.1.0.2). Data visualisation was performed using Matplotlib^73^ and Seaborn (v.0.11.2). Analysis and scripting were performed in Conda environments for package management. GitLab was used for code base management.

### Data availability

All of the data analysed in the manuscript are publicly available. Whole slide images (WSIs) are available from BioStudies: S-BSST383 ^44^ and S-BSST384 ^45^. Whole genome sequencing FASTQ files are available in the European Nucleotide Archive: PRJEB37808 ^44^ and PRJEB15138 ^45^.

## Results

### Using artificial intelligence to predict tumour genetics from digital pathology

We sought to develop a deep transfer learning model, validated by machine learning frameworks, that uses clinical grade H&E whole slide images (WSIs) to predict germline and somatic genetic alterations. Our framework leveraged data from 597 experimentally-controlled, DEN (N-diethylnitrosamine) induced liver tumours from four mouse strains (**Fig. 1A**). Critically, each WSI was associated with whole-genome sequencing data derived from the immediately adjacent fresh tumour sample, which we recently characterised^44,45^. Therefore, tumour-specific germline variation and somatic alterations were available as ground-truth for training the prediction models.

Tumour WSIs were annotated and tessellated to fixed-size non-overlapping 256 x 256 µm tiles within the QuPath^53^ environment (total tiles=163,835; mean tiles per tumour=274; **Fig. 1A**). Next, we followed two technically distinct computational pathology strategies for phenotype-to-genotype prediction: (i) a weakly supervised deep transfer learning pipeline in which the algorithm was permitted to discover salient image features, and (ii) a supervised machine learning pipeline based on quantification of human-interpretable features of nuclear morphology.

In the first approach, we applied weakly supervised deep transfer learning pipelines based on Convolutional Neural Networks (CNNs) (MobileNetV2, ResNet50, InceptionV3, VGG19) to the 163,835 H&E image tiles derived from 597 tumour WSIs to make slide-level predictions of (i) germline variation, (ii) somatic driver mutation, and (iii) genome (a)symmetry (**Fig. 1B; Methods**). These four base CNN models were initially trained using ImageNet weights, and then fine-tuned to our dataset. For the second approach, seeking to add human interpretability, we identified and segmented 22,879,653 epithelioid tumour nuclei in the H&E images using StarDist (Schmidt et al. 2018) within QuPath (segmentation precision: median=0.880, range: 0.548-0.990; detection probability: median=0.758, range: 0.697-0.807; **Supplementary Fig. 1B,C**; **Methods**). For each nucleus, we quantified five metrics of nuclear geometry (area, perimeter, circularity, maximum diameter, and minimum diameter) and ten colourimetric measures of optical density (mean, median, minimum, maximum, and standard deviation; quantified separately for haematoxylin and eosin). These nucleus-level measurements were then abstracted to tile-level and WSI-level using summary statistics of both central tendency and variance. We modelled nuclear features using four machine learning approaches (Logistic Regression, Support Vector Classifier, Random Forest Classifier, Gradient Boosting Classifier) to make slide-level predictions of the same groups of genetic alterations (**Fig. 1B**; **Methods**). All eight models were trained following a 70/30 cross-validation partition strategy prior to evaluation on held-out test data (**Methods**).

For each WSI (tumour) and each classification task, we constructed a ROC (Receiver Operating Characteristic) curve from the binary classification (“correct” vs “incorrect”) of all its component tiles, yielding a single per-tumour AUC (Area Under the Curve) value (**Fig. 1B**). This approach evaluates model performance at the individual tumour level (rather than across tumours), where an AUC of 1.0 indicates perfect slide-level (i.e. tumour) classification and 0.5 implies equivalent predictive ability as random chance. In the subsequent results, ResNet50 performed the best across prediction tasks for transfer learning, and Gradient Boosting Classifier (GBC) for supervised machine learning. For clarity we focus on the comparison of these two approaches, and the performance of all considered models is evaluated in the supplementary figures.

### Genome-wide and locus-specific germline variation are resolved by H&E staining alone

Ancestral genetic variation, comprising millions or tens of millions of polymorphisms between individuals, may influence the biological attributes and clinical behaviour of human cancers^37–42^. However, investigating these effects is challenging in light of global and regional differences in environmental exposures, socioeconomic factors, and healthcare provision, in addition to systemic biases in the design and implementation of large studies. When insufficiently diverse training data are utilised, machine learning systems may be vulnerable to ancestry biases^74–76^. To test whether this genome-wide genetic variation impacts cellular morphology independently of confounding effects, we analysed a cohort of liver tumours of the same histological subtype that were experimentally induced in four strains of mice (**Fig. 2A**). These reflect similar evolutionary divergence to that present in the human population^45^.

**Fig.2.**
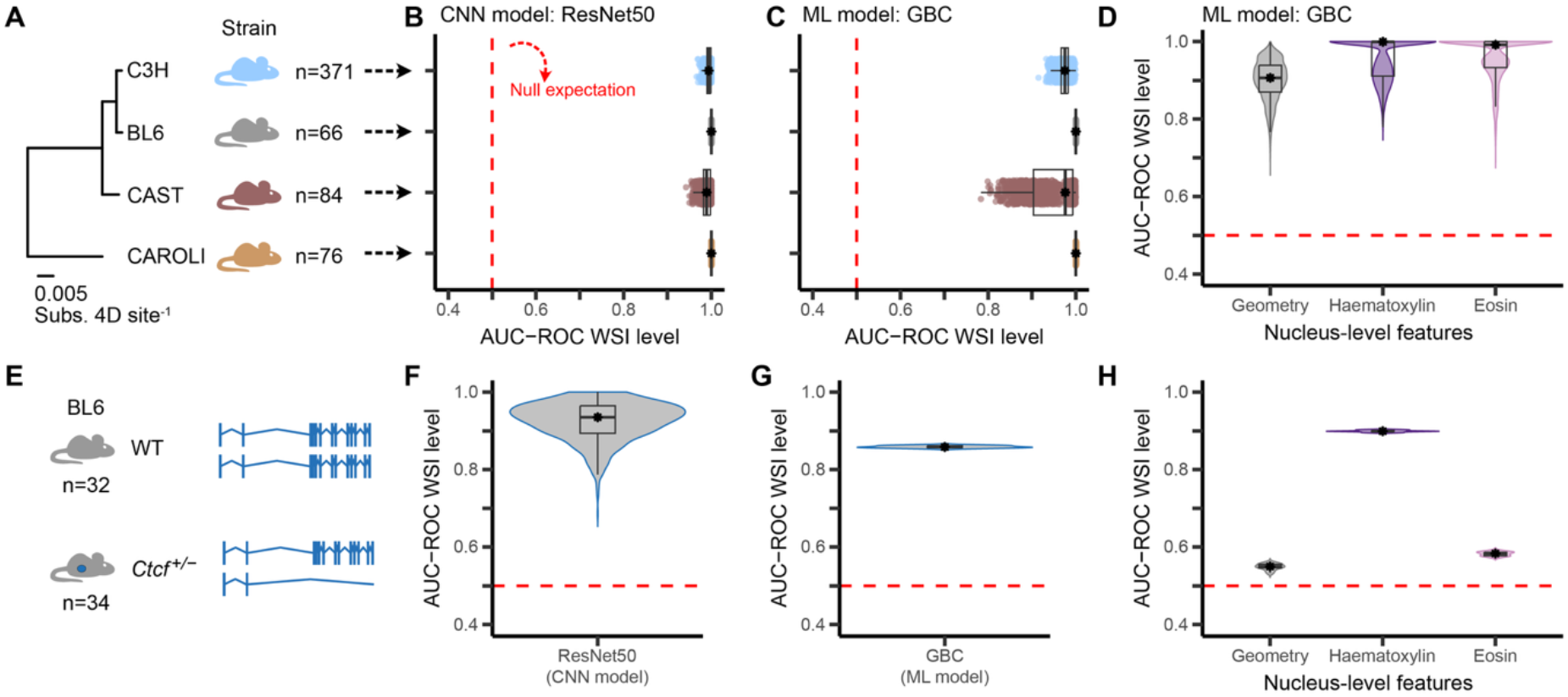
Histochemical staining intensities predict both genome-wide and locus-specific germline differences. **(A)** Summary of study cohort. Liver tumours were chemically-induced in four strains of evolutionarily divergent mice (phylogenetic tree branch lengths are substitution rates of protein coding 4-fold degenerate sites). The number of tumours per strain is indicated. **(B)** The Convolutional Neural Network (CNN) ResNet50, trained to predict strain within a transfer learning pipeline, is highly performant across all strains. Null expectation = 0.5 (red line). Each coloured point represents the slide-level AUC-ROC value of one bootstrap replicate (1,000 per tumour); strain colours from (A). Boxplot boxes indicate first quartile, median and third quartile. Whiskers indicate the first or third quartile ± 1.5 times the interquartile range (IQR). **(C)** Machine Learning (ML) using a Gradient Boosting Classifier (GBC) trained as a binary classifier to predict strain from quantitative nuclear morphometry (see Fig. 1B) also performs well across all strains. Plotted as per (B). **(D)** Comparing subsets of nuclear features used in the GBC model identifies that colourimetry is more highly predictive for strain than nuclear geometry. Violin plots show the probability density of data, boxes indicate first quartile, median and third quartile; and whiskers indicate the first or third quartile ± 1.5 times the IQR. **(E-H)** The same ML approaches were applied to liver tumours from BL6 littermates that were either wild type (WT) or hemizygous for *Ctcf* (*Ctcf* ^+/-^) in hepatocytes, following Cre-loxP excision of the *Ctcf* coding exons (exons 3 to 12; **Methods**) (**E**). In these tumours, there was high predictive power for *Ctcf* hemizygosity using both ResNet50 (**F**) and GBC (**G**); nuclear haematoxylin staining is the dominant contributing feature to the GBC model (**H**).

First, using deep transfer learning pipelines, we trained CNNs to predict strain as a multiclass classification task. This approach, in which feature representation is learnt automatically from H&E tiles, had high performance for prediction of each strain (ResNet50 AUC range across strains=0.94-1.00, **Fig. 2B**; MobileNetV2, InceptionV3, and VGG19 in **Supplementary Fig. 2A**). Motivated by these discernable differences between strains, we next took a supervised approach to determine the importance of specific image features. We found that machine learning approaches, which used pre-quantified nuclear features to make slide-level strain prediction, were also highly predictive but with more variable performance than deep learning models (Gradient Boosting Classifier (GBC) AUC range across strains=0.78-1.00, **Fig. 2C**; Random Forest Classifier, Logistic Regression, and Support Vector Classifier in **Supplementary Fig. 2B**). Comparing subsets of nuclear features used in the GBC classifier identified that, for genome-wide germline (strain) prediction, colourimetric features were more predictive than nuclear geometry. In particular, nucleus-level haematoxylin optical density had the greatest contribution to model performance (**Fig. 2D**, **Supplementary Fig. 2C**). No evident intra-batch differences were observed for histochemical staining intensities, confirming that the strain predictions were not driven by differences in data collection over time (**Supplementary Fig. 3**).

After demonstrating that image-derived features are able to accurately discriminate genome-wide polygenic germline variation, we next applied our prediction frameworks to identify locus-specific germline alterations. To do this, we compared tumours from BL6 littermates that were either wild-type (n=32) or hemizygous (n=34) for *Ctcf* (**Fig. 2E**), which is a multifunctional genome insulator protein^77^ involved in homeostasis of cancer pathways^78,79^. We hypothesised that, since Ctcf dosage controls chromatin organisation, cell-cycle regulation, and differentiation state^77^, its depletion would impact nuclear morphology, thus providing a suitable candidate to evaluate locus-specific prediction. Both models demonstrated high predictive ability for distinguishing *Ctcf* genotype (ResNet50 mean AUC=0.93 [0.69-1.00], GBC mean AUC=0.86 [0.85-0.86]; **Fig. 2F,G**; other models **Supplementary Fig. 4A,B**), with haematoxylin optical density being the dominant contributing feature (**Fig. 2H**, **Supplementary Fig. 4C**). Haematoxylin binds negatively charged nucleic acids (including DNA) and therefore the observed variation in staining intensity between *Ctcf* genotypes may reflect altered chromatin organisation^79^, in turn affecting the accessibility of the DNA phosphate backbone for haematoxylin staining.

In summary, our deep learning models are highly predictive for both genome-wide polygenic germline variation (strain) and locus-specific germline heterozygosity between littermates of an otherwise genetically-homogeneous inbred strain. The application of supervised machine learning revealed that, in both scenarios, nuclear staining was more influential than nuclear geometry.

### Explainable prediction of altered oncogenic driver gene from H&E images

Having considered the impact of germline variation, next we tested whether we could predict acquired oncogenic mutations from tumour pathology. Somatic mutations activating the MAPK pathway are common in human cancers and are tractable targets for precision oncology^80–82^. MAPK genes are also commonly, and mutually exclusively, mutated in our cohort of DEN-induced liver tumours, with *Egfr*, *Hras*, and *Braf* driver variants identified across all four mouse strains^45^. Since we have already demonstrated that locus-specific germline variation can be detected from tumour histopathology (**Fig. 2F-H**), we hypothesised that oncogenic somatic mutations could also be predicted. Additionally, the multi-strain nature of our dataset enabled us to test the generalisability of such predictions across ancestrally-divergent populations.

We initially applied our computational frameworks to predict *Egfr*, *Hras*, and *Braf* mutations in our large cohort of C3H liver tumours (n=371 WSIs; **Fig. 3A; Supplementary Fig. 5A,B**). Restricting model training and testing to a single strain ensured that its prediction was not confounded by known strain differences in driver mutation distribution^45^. The weakly-supervised deep transfer learning pipeline using ResNet50 successfully predicted driver mutation identity (*Egfr*, *Hras*, or *Braf*) as a multiclass classification task (mean AUC across drivers=0.77 [0.44-1.00]; **Fig. 3B**). Machine learning using the GBC achieved slightly lower overall performance than the ResNet50 model (mean AUC across drivers=0.63 [0.39-0.94]; **Fig. 3C**). For somatic mutation prediction we found that, unlike germline prediction, nuclear geometric features outperformed staining features (geometry median AUC 0.60, compared with 0.56 and 0.57 for haematoxylin and eosin, respectively; **Fig. 3D**). Specifically, Shapley analysis revealed that driver prediction performance was primarily driven by measures capturing nuclear size and shape variance, rather than circularity (**Supplementary Fig. 5C**). This suggests that image-level variability in nuclear morphology represents a phenotypic link to underlying genetic alterations.

**Fig.3.**
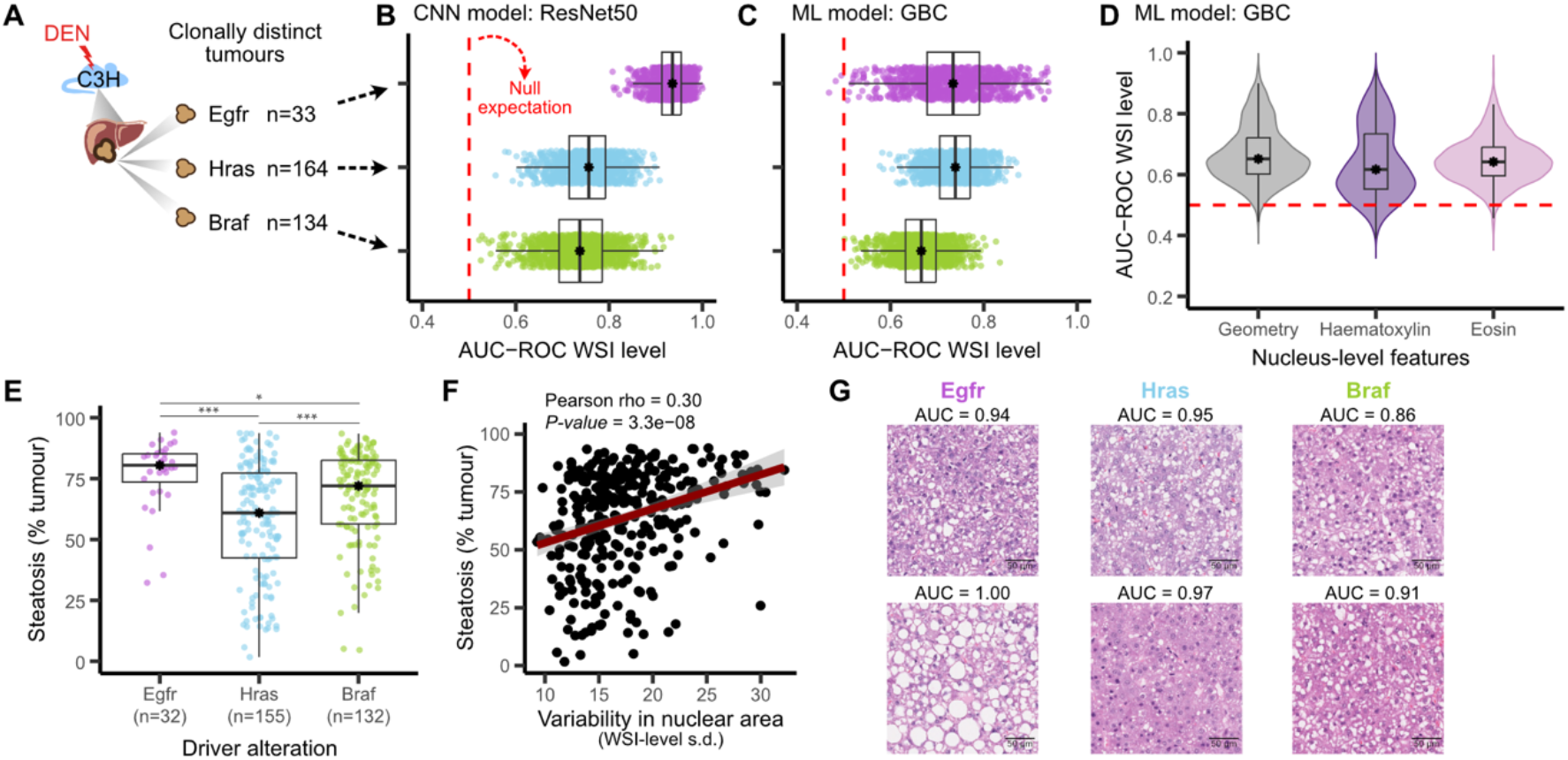
Driver mutations are associated with nuclear morphometry and variation in hepatic steatosis in C3H. **(A)** Liver tumours from C3H mice with an oncogenic mutation in a MAPK gene (*Efgr, Hras, Braf*; n=331) were used for driver mutation prediction. **(B)** Deep transfer learning using ResNet50 trained as multi-class classifiers to predict oncogenic mutations performs well for each driver mutation. *Egfr:* median AUC=0.93, *Hras*: median AUC=0.75, *Braf*: median AUC=0.74. Null expectation = 0.5 (red line). Dots represent slide-level AUC-ROC values of one bootstrap replicate (1,000 per tumour). Boxplot boxes indicate first quartile, median and third quartile. Whiskers indicate the first or third quartile ± 1.5 times the IQR. **(C)** Oncogenic driver mutations can also be predicted from quantitative nuclear morphometry using statistical machine learning (GBC). *Egfr:* median AUC=0.73, *Hras*: median AUC=0.74, *Braf*: median AUC=0.67 (1,000 bootstraps per tumour). Plotted as per (B). **(D)** Comparing subsets of nuclear features used in the GBC model identifies that nuclear geometry is more predictive for strain than optical density of H&E staining. Violin plots show the probability density of data, boxes indicate first quartile, median and third quartile; and whiskers indicate the first or third quartile ± 1.5 times the IQR. **(E)** Quantitative image analysis identified the highest amount of steatosis (percentage of tissue area) in tumours with *Egfr* driver mutations and the lowest amount in those with *Hras* mutations. Some tumour slides from (A) were excluded following image quality control (n per driver mutation indicated on the x-axis). Boxplot boxes indicate first quartile, median and third quartile. Whiskers indicate the first or third quartile ± 1.5 times the IQR; two-tailed Mann-Whitney-Wilcoxon test, * p<0.05, *** p<0.001. Bonferroni-corrected p-values are shown. **(F)** There is a significant positive relationship between the variability in nuclear area and the amount of steatosis (slide-level standard deviation of nuclear area). Scatter plot with linear fit (red) and 95% confidence interval (grey shading). **(G)** Representative high-confidence tiles from true-positive slides within the held out test set for each class show the majority of H&E tiles exhibit overlapping morphological features, regardless of driver mutation (top row). Extreme outliers show divergent steatosis phenotypes (bottom row), reflecting the quantitative trends identified from image analysis (**E,F**). Slide-level AUC-ROC is given for each tumour.

Differences in nuclear morphology may arise from either intranuclear processes, such as cell cycle phase^83^ and genome content^84^, or extranuclear factors, such as cytoplasmic lipid or glycogen inclusions that can obscure and deform nuclear cross-sectional shape^85^. To explore the biological basis of these predictive histological signals, high-confidence, true-positive tiles were manually inspected for recurrent interpretable patterns. While most tiles had very similar histopathological features, the subset of highest-confidence tiles revealed unexpected variation in tissue phenotypes between *Egfr*, *Hras*, and *Braf* mutated tumours. Specifically, histopathological review identified that both the quantity and pattern of hepatic steatosis (fat accumulation within the cell cytoplasm) varied between tumours with different driver mutations (**Fig. 3E-G**).

To formally assess these observations, we used image analysis software (Visiopharm) to segment and quantify cytoplasmic lipid droplets in each tissue slide. We found the proportion of steatotic tumour area was higher in *Braf*- and *Egfr*-mutated tumours compared to those with *Hras* mutations (Bonferroni-corrected p=4.249×10^-05^ and 6.369×10^-04^ respectively, two-tailed Mann-Whitney U tests; **Fig. 3E**), and slide-level steatosis percentage was significantly associated with nuclear morphology variability (Spearman ⍴=0.30, p=3.62×10^-8^, **Fig. 3F**). In tiles with a higher proportion of macrovesicular steatosis, nuclei were placed eccentrically within the cell and deformed to a more angular contour by the adjacent lipid droplet (**Fig. 3G**, lower left panel), whereas in cells with fewer or smaller inclusions (microvesicular) nuclei tended to be centrally placed and round in contour (**Fig. 3G**).

To assess the generalisability of these C3H-trained models, we next applied them to predict driver alterations in tumours from the other mouse strains. Although the predictive performance was reduced, the models show a moderate ability to discriminate somatic driver alterations in non-C3H tumours (ResNet50: mean AUC=0.74; GBC: mean AUC=0.54; **Supplementary Fig. 5D**). This reduction in prediction ability is likely due to the detectable germline effects identified above (**Fig. 2**), highlighting the importance of training on genetically-diverse cohorts.

The finding that specific MAPK driver mutations cause detectable changes in cellular phenotypes (e.g. cytoplasmic lipid accumulation) builds on our previous gene expression analyses. Although MAPK pathway gene expression was broadly perturbed across all tumours, *Egfr*-driven tumours show a distinct transcriptional profile compared with *Hras*- and *Braf*-driven tumours^45^. These observations provide a mechanistic bridge between oncogenic driver mutation and histology by showing that the functional output of the mutation alters cellular metabolism, and resulting composition changes distort nuclear morphology.

### Computational pathology explains strain-specific mutational symmetry

The vast majority of DEN-induced tumours in our cohort have genome-wide chromosome-scale mutational strand asymmetry as consequence of DNA lesion segregation. For example, a 90-times higher rate of T→C mutations than the reverse complement A→G mutations with respect to the forward DNA strand, across an entire chromosome. Following a single DNA-damaging exposure, replication over unrepaired damage results in mutations in the newly synthesised DNA. Each daughter cell randomly inherits one of the sister chromosomes that harbour mutations arising from damage on either the initial forward strand or the initial reverse strand^44^. As such, tumours that clonally expand from a damaged cell should have chromosome-scale strand asymmetry of mutations, reflecting the originally-damaged DNA strand (**Fig. 4A**, top panel). However, a minority of tumours had near perfect mutational symmetry (i.e. absence of the expected asymmetry)^44,45^, which occurred significantly more frequently in CAROLI mice than the other strains (38.16% [CAROLI] versus 0-3% [C3H, BL6, CAST], p=2.46×10^−18^, Fisher’s exact test; **Fig. 4B**).

**Fig.4.**
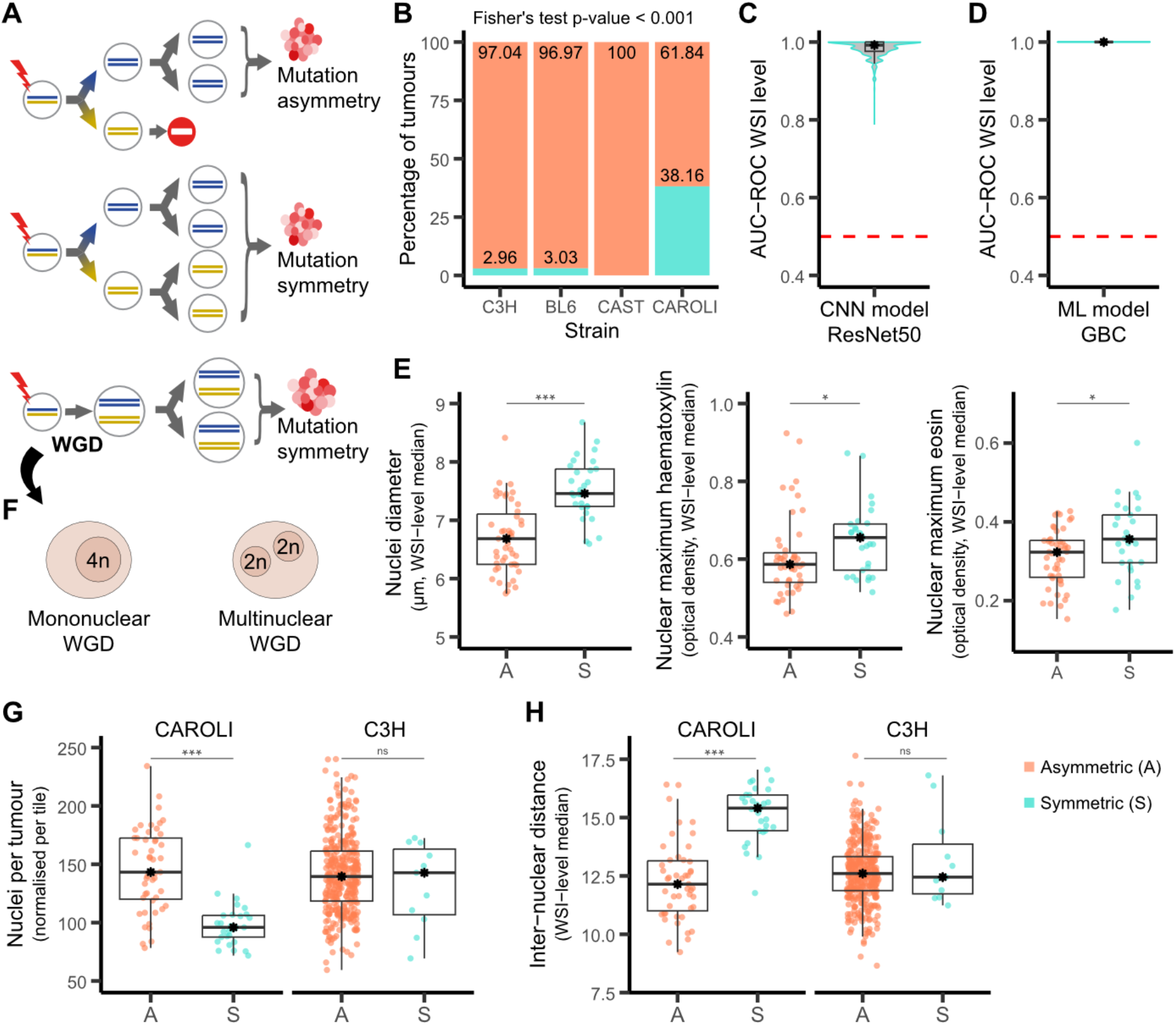
Morphometry-based models are highly performant in predicting genome symmetry in CAROLI and are driven by nuclear size. (**A**) Following an episode of damage, DNA lesion segregation ^44^ causes chromosome-scale, genome-wide asymmetry of mutations (top panel). However, mutation symmetry can occur if there is survival and expansion of both mirrored ‘sister twins’ (middle), or due to whole genome duplication (WGD, lower panel). (**B**) The vast majority of C3H, BL6, and CAST tumours have mutationally asymmetric genomes (orange), whereas over a third (n=29/76) of CAROLI tumours have mutation symmetry (green). (**C**) The weakly supervised deep transfer learning pipeline trained using ResNet50 successfully predicts genome symmetry in CAROLI tumours (slide-level AUC-ROC median=0.99; 1,000 bootstraps per tumour). Violin plots show the probability density of data, boxes indicate first quartile, median and third quartile; and whiskers indicate the first or third quartile ± 1.5 times the IQR. (**D**) The machine learning algorithm (GBC) based on quantified nuclear features also predicts genome symmetry (slide-level AUC-ROC median=1.00; 1,000 bootstraps per tumour). Violin plots show the probability density of data, boxes indicate first quartile, median and third quartile; and whiskers indicate the first or third quartile ± 1.5 times the IQR. (**E**) Nuclei in mutationally symmetric (S) CAROLI tumours are larger (short-axis length, left) than asymmetric (A) tumours (p=6.21×10^-8^, two-tailed Mann-Whitney-Wilcoxon test). Symmetric tumours also show increased histochemical staining with haematoxylin (middle panel; p=1.37×10^-2^, two-tailed Mann-Whitney-Wilcoxon test) and eosin (right panel; p=3.19×10^-2^, two-tailed Mann-Whitney-Wilcoxon test). (**F**) Schematic showing that mitotic failure (the most common cause of WGD in human cancers^97,105^) can be caused by either (i) failure of karyokinesis (nuclear division), resulting in progeny with a duplicated tetraploid genome contained within a single nucleus per cell (mononuclear WGD, i.e. 4n); or (ii) failure of cytokinesis (cytoplasmic division) alone, resulting in progeny with two nuclei within a single cell (multinuclear WGD, i.e. 2n + 2n). In either scenario, total cell size is expected to increase proportionally with polyploidy^87,88^, but the nuclear size and spacing would be different. (**G**) Tumours with mutation symmetry have fewer nuclei per tile than asymmetric tumours in CAROLI (p=1.59×10^-9^, two-tailed Mann-Whitney-Wilcoxon test) but not in C3H mice (p=0.55, two-tailed Mann-Whitney-Wilcoxon test). (**H**) The nuclei of tumours with mutation symmetry were spaced further apart than asymmetric tumours in CAROLI (p=7.51×10^-11^, two-tailed Mann-Whitney-Wilcoxon test) but not in C3H mice (p=0.73, two-tailed Mann-Whitney-Wilcoxon test).

Mutationally symmetric tumour genomes can arise under at least two different biological scenarios: First, when whole genome duplication (WGD) occurs in the first cell cycle following the mutagenesis event^45^ (**Fig. 4A**, lower panel); or second, when both daughters of the mutagenised cell persist and clonally expand together^86^ (**Fig. 4A**, middle panel). Both of these scenarios mask the mutation asymmetry of lesion segregation, resulting in perfectly (or near perfectly) symmetric genomes (**Fig. 4A**). We aimed to distinguish between asymmetric- and symmetric-genome tumours from digitised histopathology images, and determine the underlying aetiology of mutation strand symmetry.

We trained our deep transfer learning and machine learning algorithms to predict genome symmetry in CAROLI tumours classified as having symmetric (n=29) or asymmetric genomes (n=47), achieving robust prediction performance with both approaches (**Fig. 4C,D**, **Supplementary Fig. 6A,B**). This was predominately driven by the central tendency of nuclear size (**Supplementary Fig. 6C**), which were significantly larger in symmetric-genome tumours (short axis, median per slide, p=6.21×10^-8^, two-tailed Mann-Whitney-Wilcoxon test; **Fig. 4E**; also reported in^45^). The maximum optical density of nuclear haematoxylin and eosin staining (median per slide) was also significantly increased in symmetric-compared to asymmetric-genome CAROLI tumours (p=1.37×10^-2^ and p=3.19×10^-2^, two-tailed Mann-Whitney-Wilcoxon test; **Fig. 4E**). Taken together, increased nuclear size and increased chromatin staining in symmetric-genome CAROLI tumours are consistent with an overall increase in genome content, supporting that these tumours have undergone WGD. In contrast, when these models trained in CAROLI were generalised to tumours arising in C3H mice (n=11), their performance depreciated significantly (**Supplementary Fig. 6D**) which could reflect strain effects on nuclear morphology (as for driver prediction) or distinct WGD mechanisms.

This framework for WGD prediction is based on the finding of larger nuclei in CAROLI symmetric-genome tumours, and therefore assumes mononuclear WGD (i.e. a single 4n nucleus per cell, **Fig. 4F**). Multi-nuclear WGD (e.g. two or more 2n nuclei per cell, **Fig. 4F**) tumours might not be reliably identified by this approach since their nuclei would have similar morphology to diploid cells and simply be more numerous^87,88^. We investigated whether differences in polyploidisation (mononuclear vs multinuclear) accounted for depreciation of symmetric-genome model prediction in C3H tumours by comparing the distribution and density of nuclei (number of nuclei per tile) between tumours.

In CAROLI tumours, we observed a significant reduction in the number of nuclei per tumour area in those with symmetric-genomes, indicative of larger cells; in contrast, no significant difference was observed in C3H tumours (**Fig. 4G**). In addition, symmetric-genome CAROLI tumours showed increased inter-nuclear distances to their nearest-neighbour nuclei compared to asymmetric tumours; this difference was not found in symmetric C3H tumours (**Fig. 4H**). These observations further support a process of mononuclear WGD in symmetric CAROLI tumours (**Fig. 4A**, middle panel), but not in symmetric C3H tumours. Indeed, together with parallel analysis of variant allele frequency distributions^45^, nucleotide excision repair^86^, and mitochondrial content^89^, these observations suggest that these rare symmetric-genome C3H tumours most likely arise from the persistence and clonal expansion of both diploid daughter cells (**Fig. 4A**, middle panel), rather than from tetraploidisation (**Fig. 4A**, lower panel).

## Discussion

Digital pathology, cancer genomics, and artificial intelligence (AI) are defining a new era of clinical biomarkers^30,90^. This proof-of-concept study relates histologic to genomic pathology in a large cohort of experimentally-induced liver tumours, integrating contemporary computer vision methods and machine learning algorithms. All tumours in our study had the same histopathological diagnosis and near-identical cellular morphology, which creates a more computationally challenging task than deep learning studies that categorise tumours of with distinct histological appearances (e.g. lung adenocarcinoma vs squamous cell carcinoma^20,91^, or colorectal cancers with, versus without, microsatellite instability^92,93^). Despite this, we demonstrate that our deep learning models have high predictive performance and, combined with orthogonal supervised machine learning models, yield biologically meaningful and human-interpretable insights across distinct prediction tasks.

Regardless of somatic driver mutation, the majority of high-confidence tiles from true-positive slides showed overlapping, homogenous morphological features. And yet, deep learning models accurately predicted driver mutation status across ∼400 C3H tumours. Supervised machine-learning models revealed that this high prediction performance was mainly driven by differences in nuclear geometric features, particularly the variation in nuclear size and shape. Expert review of extreme outlier tiles identified divergent steatosis phenotypes, which was formalised by quantification of lipid droplets in H&E WSIs. The altered nuclear morphology, due to external compression by cytoplasmic vacuoles, offers a human-interpretable explanation for the driver prediction model. Tumours harbouring an *Efgr* driver mutation showed the highest degree of steatosis, consistent with the known role of Egfr in lipid metabolism and steatosis^94,95^, which provides orthogonal biological support for our model explainability. Our discovery may help to explain how neural networks can make predictions exploiting subtle changes in cellular morphology across images that are ostensibly ‘invisible’ to pathologists.

The power of integrative analysis was also demonstrated in making genome symmetry predictions. Based on the computational pathology image-based analyses presented here, as well as orthogonal genome analyses^45,86^, we identify that symmetric CAROLI tumours have mononuclear whole genome duplication (WGD), whereas rarer symmetric tumours in other strains are composed of ‘twin sister’ diploid clones. WGD is a clinically relevant macro-evolutionary event which allows cells to tolerate aneuploidy and genomic instability^96^. WGD is prevalent across human cancers^97^ and, despite being correlated with poor prognosis^97^, confers physiological vulnerabilities^98^ making it a tantalising target for precision oncology.

Differences in genetic ancestry have been shown not only to impact the incidence and prognosis of some cancer types, but also the pattern of somatic mutations present^39^. Our well-powered experimental model system controls biological and environmental biases that can confound these morpho-molecular association studies in humans^41,99,100^, and excludes technical differences^101^ that could mimic genetic effects. We show that germline variation (both genome-wide and locus-specific) contributes to distinctive cellular phenotypes in tumours with essentially identical classification by a pathologist. As such, models trained to predict somatic mutations in one germline background do not generalise well to other ancestries. Additionally, model optimisation was constrained in our cohort due imbalanced sample distribution across strains: 371 of the 597 analysed tumours were derived from C3H mice, yielding too few tumours per driver per genetic background for separate training in non-C3H strains. This was true both for end-to-end CNN approaches and nuclei-based models. We found that germline differences, unlike somatic alterations, resulted in differences in colourimetric staining intensities; for example, altered chromatin organisation in *Ctcf* hemizygosity impacts the optical density of haematoxylin staining. Additionally, prediction of genome symmetry in non-CAROLI tumours was poor because we found the underlying biological mechanisms differ between strains: in CAROLI there is mononuclear WGD, whereas in other strains it reflects twin sister clones.

By relating histological to genomic pathology, we have shown that genetic information can be inferred from H&E images alone, creating opportunities to integrate morpho-molecular inference into routine tissue pathways and morphology-based diagnostics. Such tools could reduce manual workload and support the assessment of challenging or borderline cases, particularly tumours with a broad differential diagnosis, such as HCC versus other low grade proliferative lesions, which rely on molecular testing^102,103^. However, performance may not generalise to samples whose genetic backgrounds lie outside the diversity represented in the training data. Algorithm design and training should adequately sample the full range of ancestral variation in the target population. As computational and genomic pathology increasingly inform precision medicine, rigorous evaluation on complex, ground truth datasets become an important but challenging frontier^104^. We show that a well-controlled *in vivo* model system with multiple complementary measurements can provide a robust and flexible environment for the evaluation of competing machine learning approaches.

## Acknowledgements and funding

We thank the CRUK Cambridge Institute Histopathology and ISH Core Facility for generating digitised whole slide images: (C. Brodie, L.-A. McDuffus, and J. Arnold); the MRC Toxicology Unit Histopathology Facility for provision of Visiopharm software (M. Southwood, C. Ficken); and members of the Aitken and Taylor labs for feedback on the manuscript.

This work was supported by Cancer Research UK (Cambridge Institute core award 20412, 22398 and strategic award 22398), MRC Human Genetics Unit core funding programme grants (MC_UU_00007/11, MC_UU_00007/16, MC_UU_00035/1, MC_UU_00035/2), MRC Toxicology Unit core funding programme grants (RG94521 and MC_PC_24012), and European Molecular Biology Laboratory core funding. This work made use of resources provided by the Edinburgh Compute and Data Facility (ECDF), including the IGC_Eddie3 high performance storage arrays (MC_PC-MR/X013677/1). J.C. was supported by a Wellcome Trust PhD Training Fellowship for Clinicians (WT223088/Z/21/Z) as part of the Edinburgh Clinical Academic Track (ECAT) programme, and a pump priming award from Edinburgh Pathology. B.H. is supported by a La Caxia Junior Leader Fellowship (ID 100010434; LCF/BQ/PR23/11980033) and is hosted by the Centro Nacional de Investigaciones Oncológicas (CNIO), which is supported by the Instituto de Salud Carlos III and recognized as a ‘Severo Ochoa’ Centre of Excellence (ref. CEX2024-001442-S) by the Spanish Ministry of Science and Innovation (MCIN/AEI/ 10.13039/501100011033). S.J.A. received a Wellcome Trust PhD Training Fellowship for Clinicians (WT106563/Z/14/Z and WT106563/Z/14/A), National Institute for Health Research (NIHR) Clinical Lectureship, CRUK Clinician Scientist Fellowship (RCCCSF-May23/100001), and support from The Pew Charitable Trusts and The Alexander and Margaret Stewart Trust.

For the purpose of open access, the authors have applied a Creative Commons Attribution (CC BY) licence to any Author Accepted Manuscript version arising from this submission.

## Author contributions

J.C. - conceptualisation, method development, formal analysis, visualisation, and writing original manuscript. B.H. - formal analysis, visualisation, and writing original manuscript. J.L. - genomic analysis of mutation symmetry. C.J.A. and S.A. - supporting analyses. P.B. - supervision of image processing. F.C. - provided tissue resources. P.F. - computational infrastructure support. D.T.O. - access to unpublished data and funding. M.S.T. - conceptualisation, formal analysis, visualisation, writing original manuscript, supervision, and funding. S.J.A. - conceptualisation, histopathology analysis, visualisation, writing original manuscript, supervision, and funding. C.A.S., P.F., D.T.O., M.S.T., S.J.A - led the Liver Cancer Evolution Consortium. All authors had the opportunity to edit the manuscript. All authors approved the final manuscript.

## Liver Cancer Evolution Consortium members

Sarah J. Aitken, Stuart Aitken, Craig J. Anderson, Claudia Arnedo-Pac, John Connelly, Frances Connor, Maëlle Daunesse, Ruben M. Drews, Ailith Ewing, Christine Feig, Paul Flicek, Paul A. Ginno, Vera B. Kaiser, Elissavet Kentepozidou, Erika López-Arribillaga, Núria López-Bigas, Juliet Luft, Margus Lukk, Duncan T. Odom, Oriol Pich, Tim F. Rayner, Colin A. Semple, Inés Sentís, Vasavi Sundaram, Lana Talmane, Martin S. Taylor & Jan C. Verburg

---

**Supplementary Fig.1.**
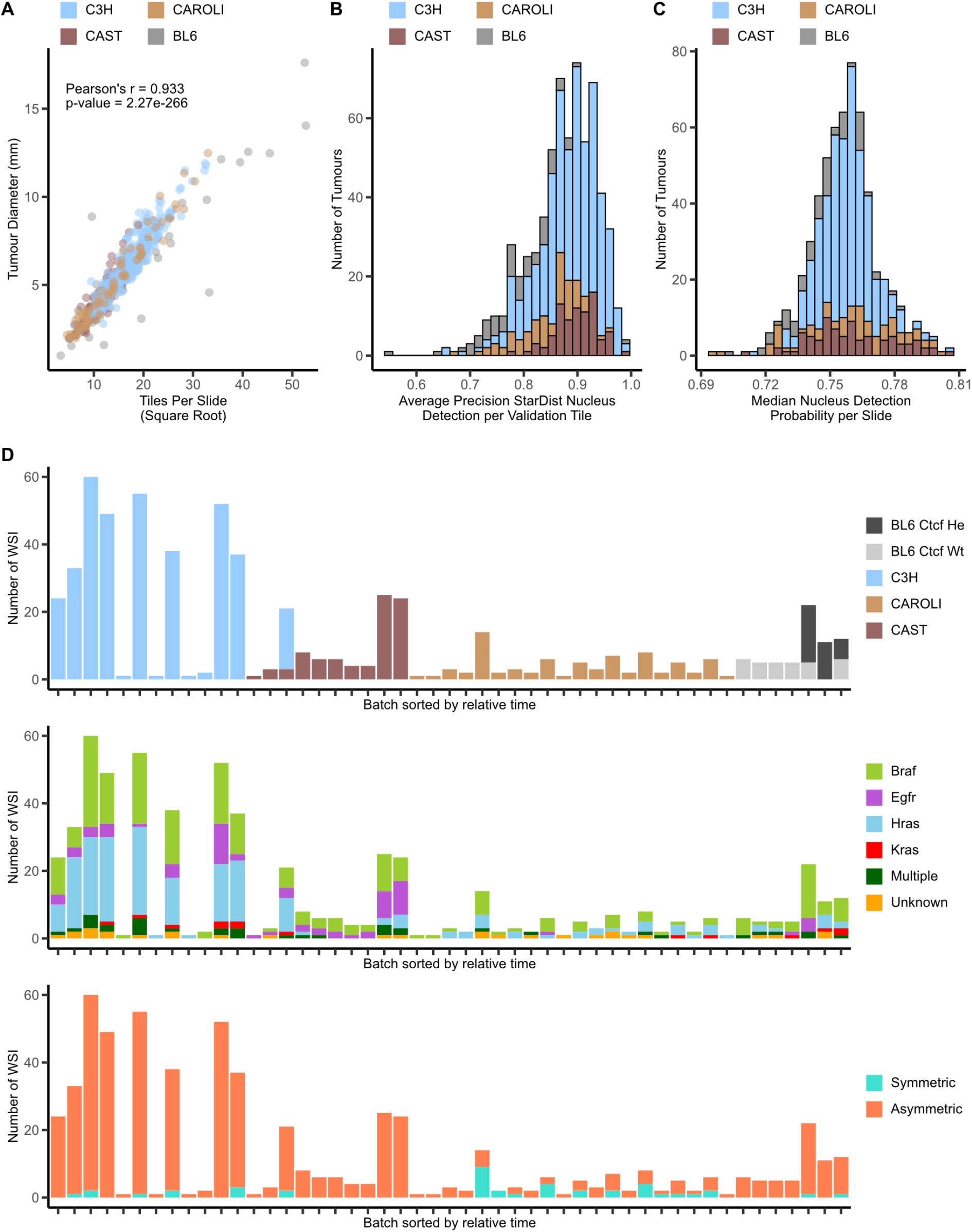
Quality metrics of tumour slides stratified by germline and somatic classification. (**A**) Scatter plot showing the association between tumour diameter (mm) and the number of tiles generated per slide (each point is one slide=tumour). (**B**) Distribution of the average precision of StarDist in segmenting epithelioid nuclei in the validation cohort, stratified by germline species. (**C**) Distribution of StarDist detection probabilities (median per slide), stratified by germline species. (**D**) Pseudotime series showing number of tumours (WSI) per processing batch stratified by germline variation (top), somatic driver oncogene mutation (middle), and genome symmetry status (bottom). Co-segregation of strains with processing batch, and genome symmetry with CAROLI strain, is observed reflecting differences in tumour latency and whole genome duplication status across strains^45^.

**Supplementary Fig.2.**
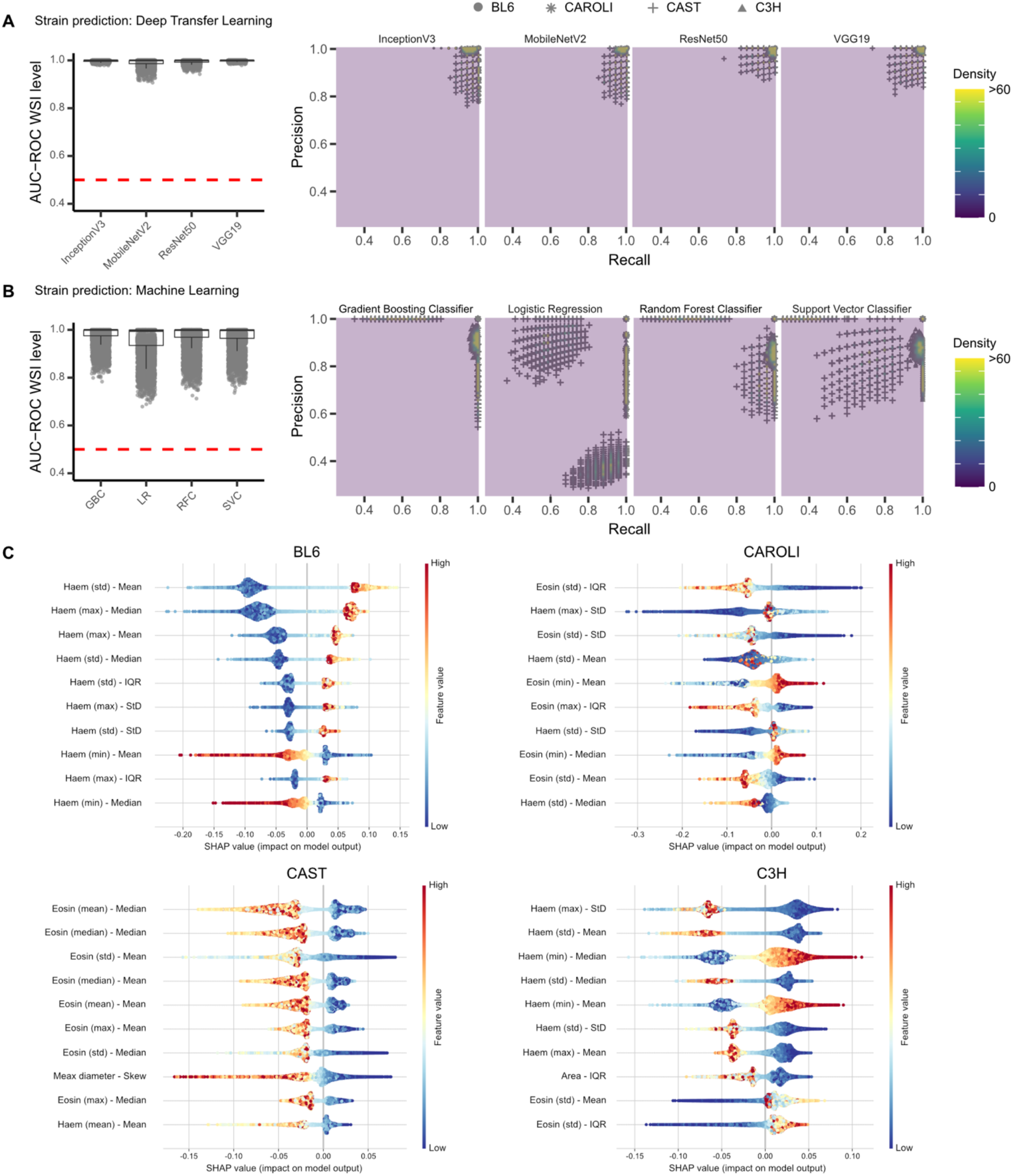
Performance of all models across strains for germline variation prediction. **(A)** Strain prediction for the four deep transfer learning pipelines based on Convolutional Neural Networks (CNNs; InceptionV3, MobileNetV2, ResNet50, VGG19). Slide-level AUC-ROC values (1,000 bootstraps per tumour) are shown for all strains together (left); null expectation = 0.5 (red line); boxplot boxes indicate first quartile, median and third quartile. Whiskers indicate the first or third quartile ± 1.5 times the interquartile range (IQR). Precision-recall performance metrics are shown (right); strain indicated by point shape. Density represents a two-dimensional kernel density estimate of event coordinates. All models have high predictive performance but more variable precision-recall. **(B)** Performance of the four machine learning approaches (Logistic Regression, Support Vector Classifier, Random Forest Classifier, Gradient Boosting Classifier) to make slide-level predictions of genetic alterations based on modelling nuclear features (1,000 bootstraps per tumour); plotted as per (**A**). **(C)** Shapley summary plots for inference of all four strains. Shapley values were computed in the held out test set for each binary random forest classifier and each point represents a tile. The top 10 tile-level features, ranked by the sum of absolute values, are plotted. Each point is colour-coded by its feature value expressed as a Z-score. Features are labelled in the format ‘feature name (nuclear level statistic where appropriate)’ - ‘tile level statistic’.

**Supplementary Fig.3.**
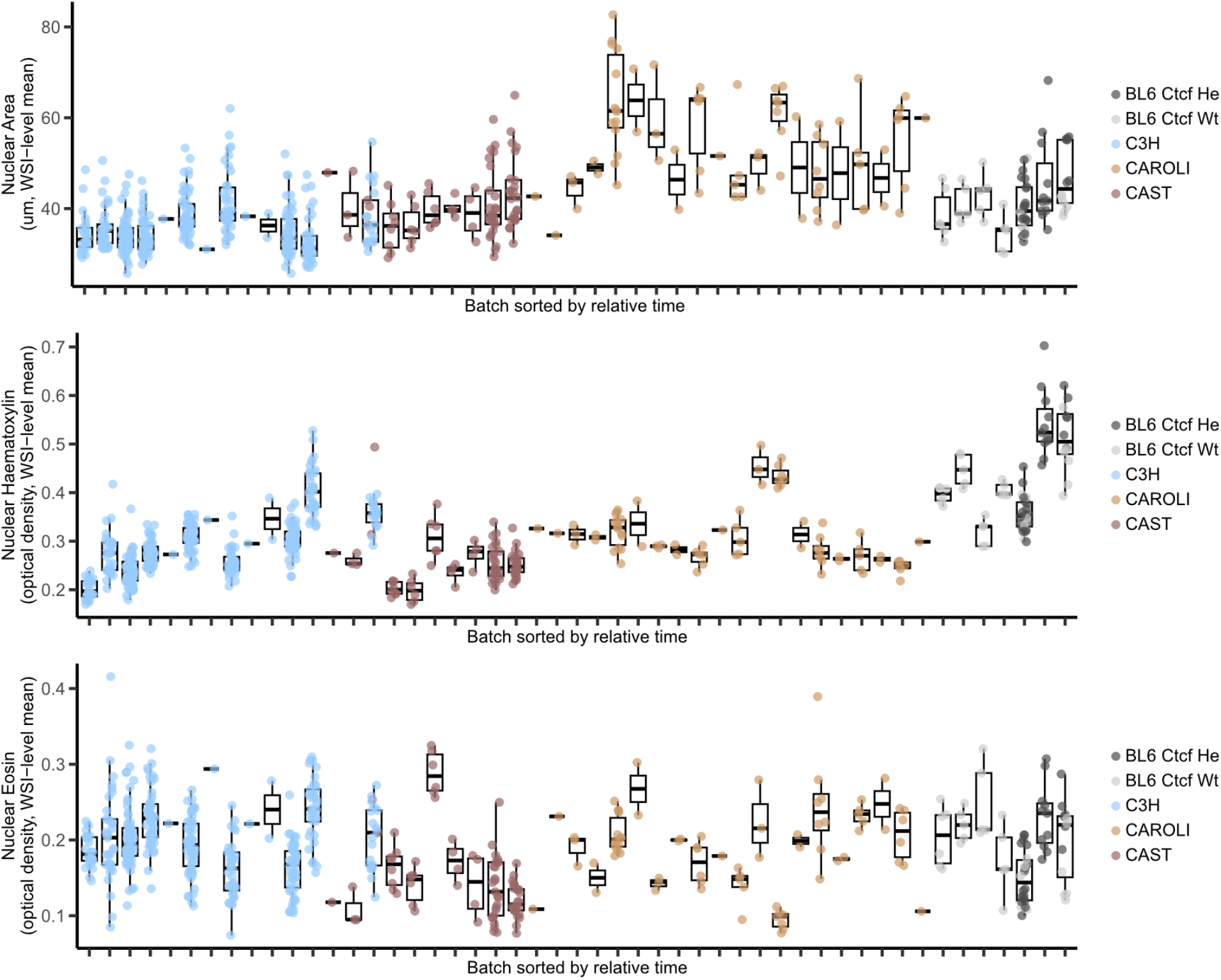
Nuclear features per sample processing batch, sorted by pseudotime. Dots represent the WSI-level mean nuclear area (top), and haematoxylin (middle) and eosin (bottom) mean staining intensities. Dots are coloured by genome-wide (strain) and locus-specific (*Ctcf* genotype, hemizygous (He) or wild-type (Wt)) germline differences. Boxplot boxes indicate first quartile, median and third quartile. Whiskers indicate the first or third quartile ± 1.5 times the interquartile range (IQR). Higher nuclear area observed in some CAROLI tumours reflects whole genome duplication (Fig. 4).

**Supplementary Fig.4.**
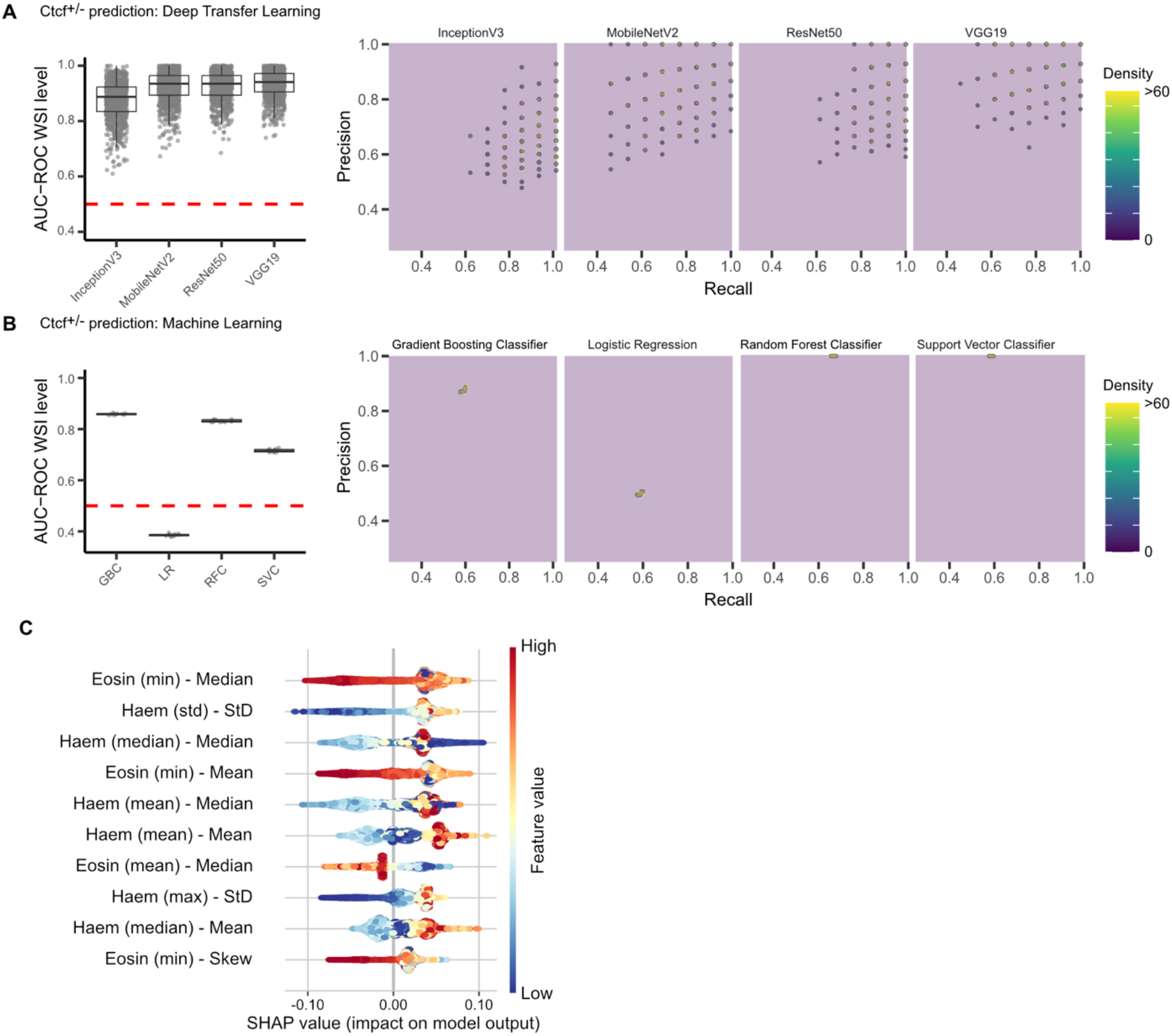
Performance of all models for locus-specific *Ctcf* genotype prediction. **(A)** Prediction of the locus-specific *Ctcf* genotype in BL6 tumours for the four deep transfer learning pipelines based on Convolutional Neural Networks (CNNs; InceptionV3, MobileNetV2, ResNet50, VGG19). Slide-level AUC-ROC values (1,000 bootstraps per tumour) are shown for all strains together (left); null expectation = 0.5 (red line); boxplot boxes indicate first quartile, median and third quartile. Whiskers indicate the first or third quartile ± 1.5 times the interquartile range (IQR). Precision-recall performance metrics are shown per strain (right). Density represents a two-dimensional kernel density estimate of event coordinates. All models have high predictive performance but more variable precision-recall. **(B)** Performance of the four machine learning approaches (Logistic Regression, Support Vector Classifier, Random Forest Classifier, Gradient Boosting Classifier) to make slide-level predictions of locus-specific genotypes based on modelling nuclear features (1,000 bootstraps per tumour); plotted as per (A). **(C)** Shapley summary plots for inference of locus-specific genotype (*Ctcf* hemizygosity). Shapley values were computed in the held out test set for each binary random forest classifier and each point represents a tile. The top 10 tile-level features, ranked by the sum of absolute values, are plotted. Each point is colour-coded by its feature value expressed as a Z-score. Features are labelled in the format ‘feature name (nuclear level statistic where appropriate)’ - ‘tile level statistic’.

**Supplementary Fig.5.**
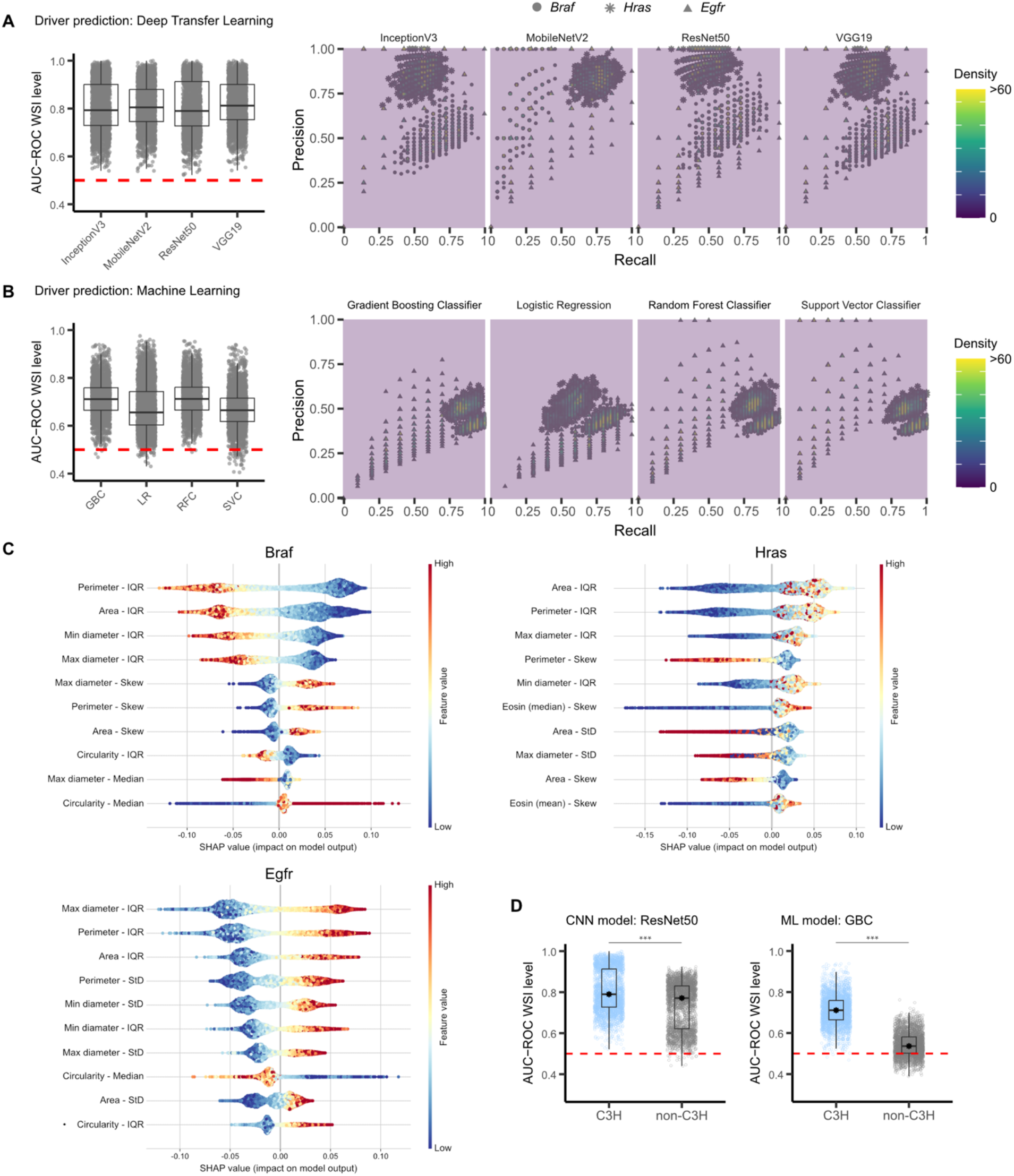
Performance of all models for driver mutation prediction. **(A)** Driver prediction in C3H tumours for the four deep transfer learning pipelines based on Convolutional Neural Networks (CNNs; InceptionV3, MobileNetV2, ResNet50, VGG19). Slide-level AUC-ROC values (1,000 bootstraps per tumour) are shown for all drivers together (left); null expectation = 0.5 (red line); boxplot boxes indicate first quartile, median and third quartile. Whiskers indicate the first or third quartile ± 1.5 times the interquartile range (IQR). Per-driver precision-recall performance metrics are shown per strain (right); driver mutation indicated by point shape. Density represents a two-dimensional kernel density estimate of event coordinates. All models have modest predictive performance and similar precision-recall. **(B)** Performance of the four machine learning approaches (Logistic Regression, Support Vector Classifier, Random Forest Classifier, Gradient Boosting Classifier) to make slide-level predictions of driver mutations in C3H tumours based on modelling nuclear features (1,000 bootstraps per tumour); plotted as per (A). **(C)** Shapley summary plots for inference of driver alterations in C3H tumours. Shapley values were computed in the held out test set for each binary random forest classifier and each point represents a tile. The top 10 tile-level features, ranked by the sum of absolute values, are plotted. Each point is colour-coded by its feature value expressed as a Z-score. Features are labelled in the format ‘feature name (nuclear level statistic where appropriate)’ - ‘tile level statistic’. **(D)** Performance of ResNet50 and GBC models trained on C3H tumours for predicting driver status in non-C3H tumours. Model performance is markedly reduced in non-C3H compared to C3H tumours (1,000 bootstraps per tumour). *** p<0.001; two-tailed Mann-Whitney-Wilcoxon test.

**Supplementary Fig.6.**
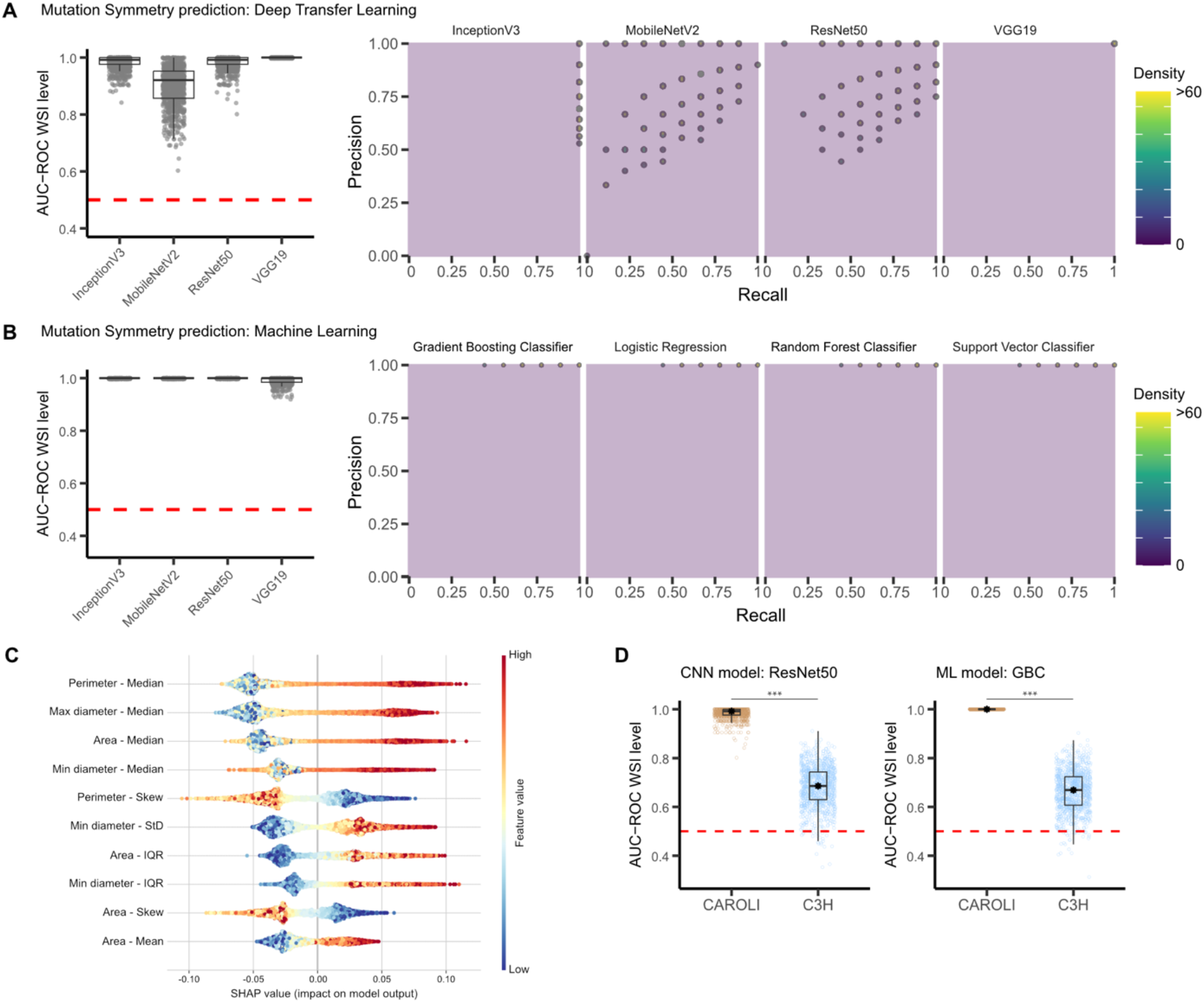
Performance of all models for mutation symmetry prediction. **(A)** Mutation symmetry prediction in CAROLI tumours for the four deep transfer learning pipelines based on Convolutional Neural Networks (CNNs; InceptionV3, MobileNetV2, ResNet50, VGG19). Slide-level AUC-ROC values (1,000 bootstraps per tumour) (left); null expectation = 0.5 (red line); boxplot boxes indicate first quartile, median and third quartile. Whiskers indicate the first or third quartile ± 1.5 times the IQR. Precision-recall performance metrics (right). Density represents a two-dimensional kernel density estimate of event coordinates. All models have high predictive performance but variable precision-recall. **(B)** Performance of the four machine learning approaches (Logistic Regression, Support Vector Classifier, Random Forest Classifier, Gradient Boosting Classifier) to make slide-level predictions of mutation symmetry in CAROLI tumours based on modelling nuclear features (1,000 bootstraps per tumour); plotted as per (**A**). **(C)** Shapley summary plots for inference of mutation symmetry prediction in CAROLI tumours. Shapley values were computed in the held out test set for each binary random forest classifier and each point represents a tile. The top 10 tile-level features, ranked by the sum of absolute values, are plotted. Each point is colour-coded by its feature value expressed as a Z-score. Features are labelled in the format ‘feature name (nuclear level statistic where appropriate)’ - ‘tile level statistic’. **(D)** Performance of ResNet50 and GBC models trained on CAROLI tumours for predicting mutation symmetry status in C3H tumours. Model performance is markedly reduced in C3H compared to CAROLI tumours. *** p<0.001; two-tailed Mann-Whitney-Wilcoxon test.

